# A neuromimetic approach to the serial acquisition, long-term storage, and selective utilization of overlapping memory engrams

**DOI:** 10.1101/621201

**Authors:** Victor Quintanar-Zilinskas

## Abstract

Biological organisms that sequentially experience multiple environments develop self-organized representations of the stimuli unique to each; moreover, these representations are retained long-term, and sometimes utilize overlapping sets of neurons. This functionality is difficult to replicate *in silico* for several reasons, such as the tradeoff between stability, which enables retention, and plasticity, which enables ongoing learning. Here, by using a network that leverages an ensemble of neuromimetic mechanisms, I successfully simulate multi-environment learning; additionally, from measurements of synapse states and stimulus recognition performance taken at multiple time points, the following network features emerge as particularly important to its operation. First, while reinforcement-driven stabilization preserves the synapses most important to the representation of each stimulus, pruning eliminates many of the rest, thereby resulting in low-noise representations. Second, in familiar environments, a low baseline rate of exploratory synapse generation balances with pruning to confer plasticity without introducing significant noise; meanwhile, in novel environments, new synapses are reinforced, reinforcement-driven spine generation promotes further exploration, and learning is hastened. Thus, reinforcement-driven spine generation allows the network to temporally separate its pursuit of pruning and plasticity objectives. Third, the permanent synapses interfere with the learning of new environments; but, stimulus competition and long-term depression mitigate this effect; and, even when weakened, the permanent synapses enable the rapid relearning of the representations to which they correspond. This exhibition of memory suppression and rapid recovery is notable because of its biological analogs, and because this biologically-viable strategy for reducing interference would not be favored by artificial objective functions unaccommodating of brief performance lapses. Together, these modeling results advance understanding of intelligent systems by demonstrating the emergence of system-level operations and naturalistic learning outcomes from component-level features, and by showcasing strategies for finessing system design tradeoffs.

## Introduction

Biological learning systems are able to acquire, store long-term, and use multiple memories whose representations overlap. For example, owl optic tecta can learn multiple associative mappings between interaural timing differences and visual field location (1), and human visual systems can adapt to input-reversing goggles and then exhibit much faster re-adaptation during subsequent reversal-goggle episodes (2). However, the storage of overlapping representations in artificial networks is notoriously elusive (3, 4). Interestingly, both brain-imitative (5) and artificial (6) input-output association systems have attempted to address this challenge using synaptic weights with two components that vary on two time scales; however, the slow weight component enables the long-term storage of at most one memory, thus falling short of natural performance. The extant literature on sensory cortex map formation, for its part, has focused mainly on how Hebbian refinement distills a promiscuously connected starting state (e.g. that observed during the critical period) into a well-patterned map, the structural specificities of which often amplify aspects of the initial condition (e.g. retinotopic projections) (7–9). It is not intuitively apparent that a network whose connectivity has annealed into a mapping, especially one favored by the initial conditions, would retain the ability to learn a second, significantly different mapping. In fact, what has been demonstrated (10) is that very early inputs can provide inertia towards certain map outcomes, but that reversible experience-driven deviations from those outcomes are possible; this learning dynamic is interestingly analogous to the single-memory long-term storage mentioned above.

Here we postulate that the mechanisms believed to govern structural plasticity could endow a system with both the flexibility to acquire multiple sensory maps and the capacity to store all of them long term. Specifically, continual synapse generation (11–13) allows brains to explore a large array of connectivity configurations. Additionally, though most exploratory synapses are pruned, those that correspond to novel environmental correlations are stabilized, and this long-term stabilization is a straightforward substrate for long-term retention. This notion is supported by observations of denser connectivity in rodents reared in enriched environments (14, 15) and of *in vivo* coincidence between long-term memory storage and long-term synapse tability (16–18). A third potential bridge between the structural plasticity literature and multi-map learning is LASG (learning-accelerated spine generation): Hebbian reinforcement, particularly of immature synapses, induces the generation of additional exploratory synapses (19–22). For an organism that learns only one map, LASG’s eventual deactivation can help account for the post-development synaptic turnover decline, while for an organism plunged into an unfamiliarly-mapped reality, LASG can perhaps disrupt the low-exploration annealed state so as to provide a learning-facilitative array of connectivity options.

In this study, we present a network that incorporates the above-described biology, and use it to model the development of mappings between multiple input modules and an output module to which they commonly project (Figure 1A, Network). In our implementation of structural plasticity, synapses initially assume the potential state, candidate synapses are generated from potential synapses, and sufficient Hebbian reinforcement can stabilize candidates permanently (Figure 1A, Synaptic states). The efficacy of existent synapses, meanwhile, is also governed by biomimetic rules: LTP, activity-dependent LTD (23, 24), and output neuron firing rate homeostasis(25) that, as in previous models of map generation (7, 8, 10), drives Hebbian competition. This network is stimulated by a series of random three-bar patterns (Figure 1A, Sample stimulus) arranged in one of four configurations (Figure 1A, Input configurations). Our simulations are composed of epochs, all bar patterns within an epoch are configured equivalently, and transitions from one epoch to the next are triggered by 1) a deceleration of new spine stabilization 2) the emergence of the output module’s ability to strongly differentiate between stimuli (Figure 1A, Test inputs) presented in the epoch’s assigned configuration. In other words, the development of response specificity to spatially transformed inputs, and of configuration-specific synapses that support said response specificity, is our study’s *in silico* analog to learning multiple sensory maps.

**Figure 1.**
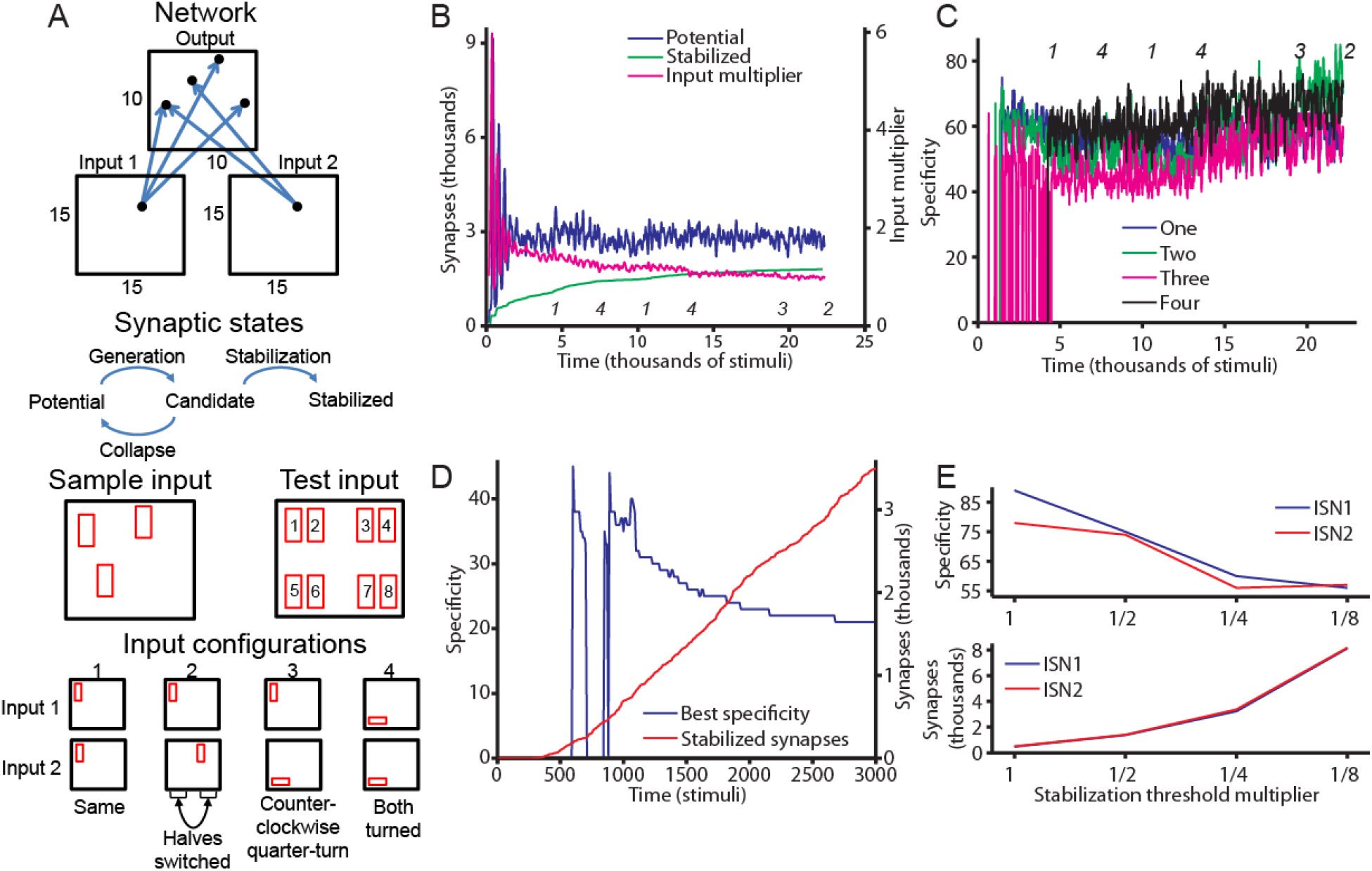
Baseline network model and its development during learning. **A** is a schematic of the circuit and modeling approach. The Network segment depicts the input and output modules, the size of their neuron grids, several neurons (black dots), and several axons (blue arrows) that connect an input neuron to a random set of output neurons, thereby serving as potential synapses. The Synaptic states segment depicts a synapse’s possible states and transitions. The Sample input segment depicts a typical stimulus presented to an input module during training. The Test inputs segment depicts the stimuli amongst which the network differentiates when response specificity is measured. Finally, the Input configurations segment depicts how each configuration affects an individual stimulus bar. **B** depicts the typical time course of connectivity development; specifically, the number of candidate and stabilized synapses, and the average homeostatic input multiplier of the output neurons. **C** depicts the specificity of the output module’s responses to the eight test stimuli from panel A. Specificity is measured across all configurations, each of which corresponds to the plotted line indicated in the legend. The depicted specificities are measured with candidate synapses removed. In panels B and C, the italicized numbers are positioned to denote epoch transitions and indicate the input configuration experienced during the preceding interval. **D** depicts, for a network whose neuronal input multipliers are fixed rather than subject to homeostatic adjustment, the number of synapses stabilized and the highest response specificity from amongst the four input configurations. Data in panels B-D generated using ISN1, and the data in B and C come from the same simulation. **E** depicts, for a series of progressively halved synapse stabilization thresholds and candidate synapse lifetimes, the response specificity and number of synapses stabilized at the endpoint of a 3000-stimulus epoch; input configuration 1 is the epoch’s input and basis for measuring response specificity.

## Results

The experiments reported below are generally replicated across two “initial state networks” (ISNs) and two distinct orderings of input configurations presented to the network. ISNs and input orderings are in turn reused across experiments, which removes these variables as confounding factors in result interpretation and reveals which experimental outcomes generalize across them. An ISN is defined as a network’s initial (randomly generated) population of potential synapses.

A full description of the model, a formal definition of response specificity, and other details are provided in *Methods*.

### Novelty causes spine generation bursts and new synapse stabilization

In our initial experiments, our network’s behaviors align with those of the biological systems that inspire its design: we witness ongoing candidate synapse turnover, and novel inputs result in the stabilization of new synapses, temporarily increased exploration of connection options, and improvement in the output module’s ability to represent the inputs. As seen in Figure 1B (green), many new synapses are stabilized during the early phases of the first two epochs, during which input configurations 1 and 4 are experienced for the first time, but the rate of new synapse stabilization is lower towards the end of these epochs and during the epochs that occur thereafter. In spite of the slowdown, novel inputs (e.g. configuration 3) continue to produce new synapses, and as the connectivity arbor gets richer, the homeostatic input multiplier (Figure 1B, magenta) declines essentially monotonically throughout the simulation. We also see in Figure 1B (blue) that the novelty experienced during the network’s initial encounters with input configurations 1, 4, and 3 produces a salient early-epoch increase in the candidate synapse count. Quantifying this effect across 4 6-epoch simulations, we find that candidate synapse count peaked during the earliest 10% of the epoch’s timespan in 11/24 epochs, which suggests a bias towards early-epoch candidate generation (one-sided binomial test, P<.001). Similarly, if we correlate epoch time with candidate synapse count, we witness (Table 1, No PING) a strong (r<-.1) and statistically significant decrease in exploratory synapse count during 13/16 of the epochs corresponding to a novel configuration, and 3/8 of those corresponding to a familiar one. Thus, the pattern of within-epoch candidate count decline is more likely to manifest (one-sided binomial test, p=.008) if the epoch’s assigned input configuration is novel. Finally, alongside the network’s connectivity, the network’s response specificity also generally develops in the expected, input-dependent manner (Figure 1C). In particular, specificity to configuration 4 was absent until inputs in configuration 4 were experienced. Meanwhile, though training with configuration 1 was sufficient to endow some response specificity to inputs in configurations 2 and 3, response specificity is markedly improved by epochs specifically devoted to these configurations.

**Table 1.** Episode-by-episode linear regression analyses of candidate count vs. time

| No PING |  |  |  | PING |  |  |  |
| --- | --- | --- | --- | --- | --- | --- | --- |
| Input | r | n | p | Input | r | n | p |
| 1<sup>†</sup> | -.1756 | 395 | < .001 | 1<sup>†</sup> | -.4389 | 382 | < .001 |
| 4<sup>†</sup> | -.2579 | 286 | < .001 | 2<sup>†</sup> | -.2963 | 286 | < .001 |
| 1 | -.0905 | 286 | .127 | 3<sup>†</sup> | -.2577 | 286 | < .001 |
| 4 | -.1809 | 286 | .002 | 4<sup>†</sup> | -.4265 | 286 | < .001 |
| 3<sup>†</sup> | -.153 | 571 | < .001 | 1 | -.0715 | 286 | .228 |
| 2<sup>†</sup> | .0125 | 286 | .833 | 3 | .0841 | 286 | .156 |
| 1<sup>†</sup> | -.178 | 353 | < .001 | 2 | .137 | 286 | .021 |
| 4<sup>†</sup> | -.2122 | 549 | < .001 | 4 | -.1958 | 286 | < .001 |
| 1 | -.1432 | 286 | .015 | 2 | .0728 | 286 | .220 |
| 4 | -.0378 | 286 | .524 | 1<sup>†</sup> | -.5028 | 420 | < .001 |
| 3<sup>†</sup> | -.1096 | 286 | .064 | 2<sup>†</sup> | .0298 | 286 | .616 |
| 2<sup>†</sup> | -.2556 | 286 | < .001 | 3<sup>†</sup> | .0478 | 286 | .420 |
| 4<sup>†</sup> | -.1735 | 439 | < .001 | 4<sup>†</sup> | -.2566 | 334 | < .001 |
| 1<sup>†</sup> | -.3111 | 333 | < .001 | 1 | .0845 | 286 | .154 |
| 4 | -.2207 | 286 | < .001 | 3 | .1125 | 286 | .058 |
| 1 | -.0851 | 286 | .151 | 2 | .0885 | 286 | .135 |
| 2<sup>†</sup> | -.2253 | 495 | < .001 | 4 | -.0111 | 286 | .852 |
| 3<sup>†</sup> | -.1519 | 286 | .010 | 2 | .1058 | 286 | .074 |
| 4<sup>†</sup> | -.2085 | 413 | < .001 | 4<sup>†</sup> | -.558 | 391 | < .001 |
| 1<sup>†</sup> | -.3327 | 286 | < .001 | 2<sup>†</sup> | -.4492 | 348 | < .001 |
| 4 | .0221 | 362 | .676 | 1<sup>†</sup> | -.1725 | 286 | .003 |
| 1 | -.0955 | 286 | .107 | 3<sup>†</sup> | -.0154 | 286 | .795 |
| 2<sup>†</sup> | -.0868 | 351 | .105 | 4 | -.1199 | 286 | .043 |
| 3<sup>†</sup> | -.2199 | 286 | < .001 | 2 | .3448 | 286 | < .001 |
|  |  |  |  | 4 | .0324 | 286 | .585 |
|  |  |  |  | 3 | .113 | 286 | .056 |
|  |  |  |  | 1 | -.0498 | 286 | .401 |
|  |  |  |  | 4<sup>†</sup> | -.5473 | 422 | < .001 |
|  |  |  |  | 2<sup>†</sup> | -.3765 | 332 | < .001 |
|  |  |  |  | 1<sup>†</sup> | -.0816 | 286 | .169 |
|  |  |  |  | 3<sup>†</sup> | -.076 | 286 | .200 |
|  |  |  |  | 4 | -.2156 | 286 | < .001 |
|  |  |  |  | 2 | .1475 | 286 | .013 |
|  |  |  |  | 4 | .1215 | 286 | .040 |
|  |  |  |  | 3 | .0785 | 286 | .185 |
|  |  |  |  | 1 | -.0768 | 286 | .195 |

Dagger superscripts (<sup>†</sup>) denote novel inputs. Correlations (Pearson's) do not quite correspond to the whole epoch; the earliest 5% is ignored in order to remove transient effects, most notably the absence of candidates at the beginning of epoch 1.

### Hebbian competition and selective correlation capture both improve response specificity

Two model features important to the above-observed emergence of tuning are homeostatic scaling and a high reinforcement threshold for synapse stabilization. We illustrate the importance of both by running ISN1 through 3000-stimulus epochs exposed to input configuration 1. Without homeostasis, the alternative is to manually set the input multiplier, which if too low renders weak nascent synapses unable to induce firing, and if set too high produces positive feedback amongst synapse stabilization events. As seen in Figure 1D, this latter scenario exhibits essentially continuous synapse stabilization with no sign of leveling off, and as more synapses are stabilized, response specificity suffers conspicuously. Meanwhile, a high reinforcement threshold acts as a low-pass filter of input correlation signals. Within short time windows, bar stimuli in different locations can correlate with each other as strongly as bar stimuli representing the same location, so stabilization rules that capture the latter will capture the former as well. Over longer periods, however, only correlations corresponding to stimulus bars in the same location (subject to configurational transformation) are encountered regularly, and over the span of a candidate synapse’s lifetime a sufficiently high reinforcement requirement can differentiate these permanent correlations from those that appear only briefly. The stabilization selectivity effects of scaling the stabilization threshold and synapse lifetime are depicted in Figure 1E.

### Inhibition-mediated filtering complements synaptic mechanisms of memory processing

Hearteningly, the above experiments indicate that our network exhibits the anticipated long-term retention ability and flexibility to represent input coincidences in seemingly any orientation and at any location. However, they also reveal a failure to switch between learned input configurations in a manner convincingly analogous to biological sensory map switches. Instead, it seems that response specificity is strong and steadily present for all input configurations once they’re learned (Figure 1C), and that the network’s stabilized synapses all receive ongoing reinforcement regardless of the input configuration being presented (Figure 2A). An even greater concern is raised by the network’s inability to learn (an example is shown in Figure 2B) all four input configurations, with either ISN, when trained with the input orderings 1234 and 4231. In part because these learning failures never manifest during a simulation’s first epoch, we attribute them to anterograde interference. Another apparent symptom of anterograde interference is that progressively fewer synapses are stabilized from one epoch to the next (Figure 1A), even though novel input configurations are being experienced.

**Figure 2.**
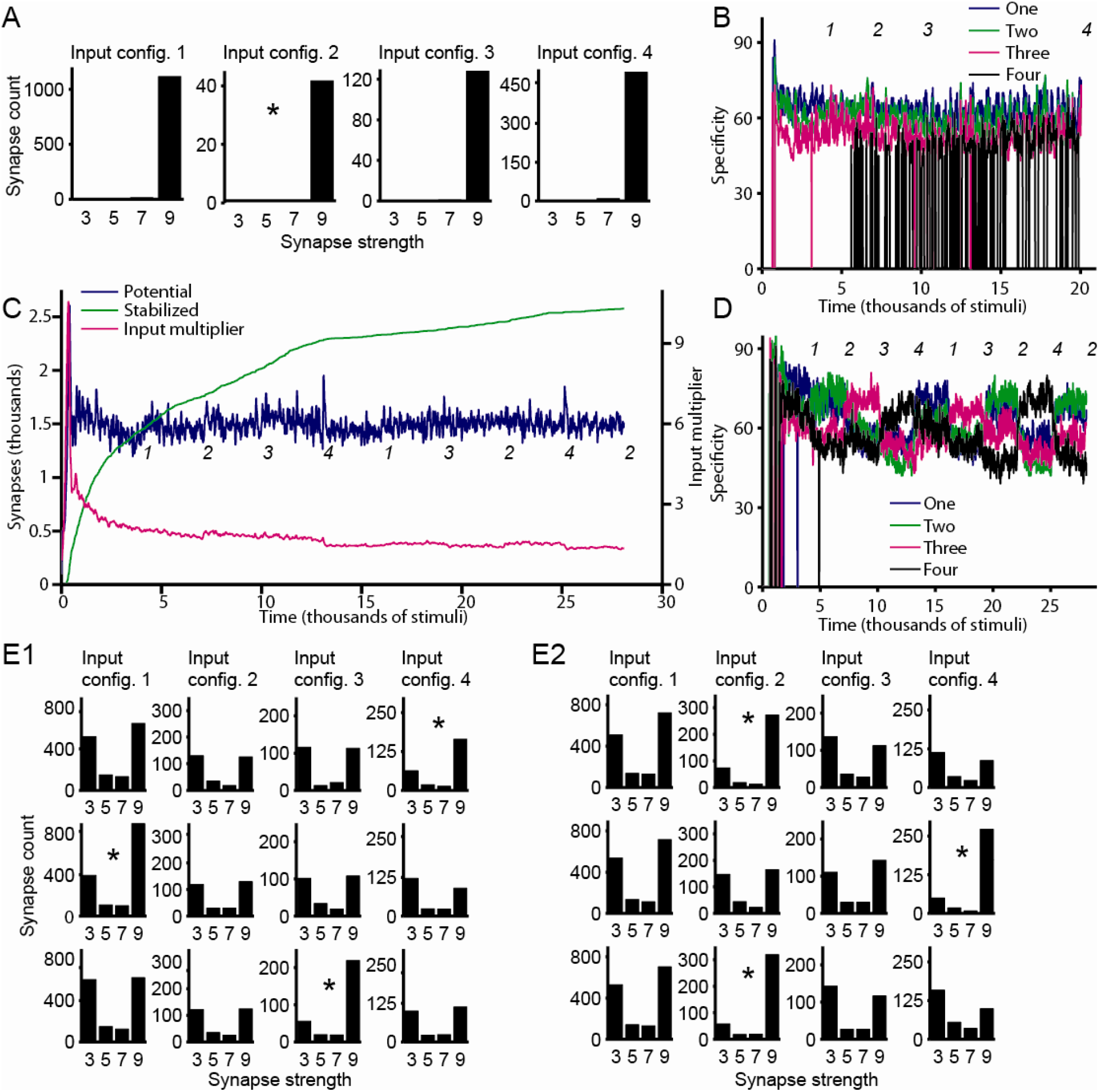
PING sharpens learning signals and suppresses passive reinforcement of presently irrelevant memory traces. **A** depicts the end-of-simulation distribution of synaptic weights, for the network whose development is depicted in Figure 1 (B and C). The four plots correspond to the synapse populations stabilized during training with each of the four input configurations. **B** depicts the development (or failure thereof) of output module response specificity in a simulation without PING and assigned the input configuration ordering 1234 (compare with Figure 1C’s 141432). **C** depicts the time course of connectivity development in a simulation with PING. **D** depicts that same simulation’s output module response specificity. Components of panels B and D (C) generally match those of Figure 1C (1B), and data are again generated using ISN1. **E1** and **E2** depict this same simulation’s epoch endpoint distribution of synaptic weights. The rows of E1 (E2) correspond to the endpoints of the middle (last) three epochs, and the four columns correspond to each input configuration’s induced synapse population. In A and E, the asterisk marks the synapse group corresponding to the epoch’s assigned input configuration.

Faced with these shortcomings, we implement an input selection mechanism analogous to pyramidal interneuronal network gamma (PING), because PING synchronizes and amplifies a network’s strongest inputs while suppressing the rest. Specifically, when the excitatory (i.e., pyramidal) neurons corresponding to the attended stimulus fire, they activate the broadly-connected inhibitory neurons, which in turn reset the network before the network’s weakly-stimulated neurons reach threshold(26). We approximate this scheme in our network simply, by adding to the output module an inhibitory cell that is reciprocally connected to all of the module’s excitatory neurons.

Just as in the no-PING experiments, the synaptic correlates of learning with PING include the coincidence of synapse stabilization with novelty (accelerated stabilization following transitions to new stimuli, and a low rate thereof during re-exposure to familiar ones, are both readily perceived; Figure 2C, green line), a downward-trending input multiplier, and elevated early-epoch candidate generation. Elaborating on the latter, we again use linear regression (Table 1, PING) to observe that within-epoch candidate counts decrease with time in 11/16 (3/20) of epochs corresponding to novel (familiar) stimuli, which suggests that this decrease behavior is strongly and preferentially associated with novelty (one-sided binomial test, P<.001). Meanwhile, candidate counts again tend to peak early; 19/36 peak during the 1^st^ 10% of the epoch (one-sided binomial test, P<.001).

Ultimately, PING delivers on our expectations that it acts to suppress activity driven by synapses from previously-learned input configurations, thereby liberating from anterograde interference the current epoch’s input and facilitating its establishment as the dominant memory trace. Specifically, we find that training the network with the input orderings 1234 (followed by 13242) and 4213 (followed by 24231) is now possible, and our simulations now exhibit epoch-to-epoch response specificity (Figure 2D) and synaptic weight (Figure 2E) shifts that emphasize each epoch’s assigned input configuration. Thanks to these shifts amongst memory traces and to the expansion of learning capability, the network’s behavior is now a much better approximation of the biology. This result is unsurprising, the role of inhibition in sensory cortex input sharpening is well-known(27, 28).

The suppression of signaling from synapses corresponding to off-epoch input configurations produces several other effects as well. One effect is input multiplier fluctuations: for example, in Figure 2C (magenta trace), the input multiplier is lower during epochs with configurations 1 and 2 than during epochs with configurations 3 and 4. We believe this is because the large synapse population corresponding to purely vertical stimulus bars (i.e., configurations 1 and 2) drives down the input multiplier when those synapses are well-reinforced, but is less effectual during other epochs. A second effect is the conspicuous (Figure 2C, blue trace) start-of-epoch candidate synapse spikes, most visible during the later epochs. Because the synapses in the no-PING simulations are never weakened and therefore cannot exhibit a salient recovery of their strength, one would expect the LASG-driven early-epoch candidate spikes to be bigger for the simulations with PING. Indeed, we find that for epochs subsequent to the first, the epoch’s peak candidate synapse count exceeds the epoch mean by three standard deviations for 19/32 of the PING epochs, while doing so far less frequently (4/20 epochs; one-sided binomial test, P<.001) in the no-PING regime.

Additional illustration of PING’s facilitation of learning comes from experiments we run without LASG (described in detail in the next section; experiment outcomes are depicted in Figures 3 and 4); in these and in the experiments described thus far, we find that simulations with PING consistently exhibit better response specificity than their counterparts without (Table 2, rows 1 and 2; comparisons performed as described in *Methods*). Moreover, all simulations producing cases of total (Figure 3, marked with “F”) or partial (Figure 3, marked with “f”) learning failure are no-PING simulations. Arguably, PING’s sharpening of learned representations generalizes existing notions of inhibition-sharpened neural signaling: neurons that fire together wire together, and specificity in the former begets specificity in the latter.

**Table 2.** Comparisons of tuning quality, two-sided rank sum tests

| Row | Conditions Compared (2<sup>nd</sup> condition better) | PING | Input Orderings | n | P | min (sig. mean diff) |
| --- | --- | --- | --- | --- | --- | --- |
| 1 | No PING vs. PING; low- and high-spine | N/A | 123413242, 432142431 | 450 (x8) | All $P < 10^{-4}$ | 8.43 |
| 2 | No PING vs. PING; LASG | N/A | 141432, 414123 | 300 (x4) | All $P < 10^{-4}$ | 12.02 |
| 3 | LASG vs. high-spine | Present | 123413242, 432142431 | 450 (x4) | All $P < 10^{-4}$ | 3.35 |
| 4 | LASG vs. low-spine | Present | 123413242, 432142431 | 450 (x4) | All $P < 10^{-4}$ | 4.29 |
| 5 | High- vs. low-spine | Present | 123413242, 432142431 | 450 (x4) | No consistent trend |  |
| 6 | LASG vs. high-spine | Absent | 141432, 414123 | 300 (x4) | All $P < 10^{-4}$ | 4.73 |
| 7 | LASG vs. low-spine | Absent | 141432, 414123 | 300 (x3), 298 | All $P < 10^{-4}$ | 5.68 |
| 8 | High- vs. low-spine | Absent | 141432, 414123 | 300 (x3), 298 | $< 10^{-4}$, .0139, .0189, .1765 | 1.53 |
| 9 | High- vs. low-spine | Absent | 123413242, 432142431 | 450 (x4) | No consistent trend |  |
| 10 | Candidates, present vs. removed | Present | All 4 | 450 (x11), 300 (x3), 449, 298 | $P < 10^{-4}$ (x16) | 2.97 |
| 11 | Candidates, present vs. removed | Absent | All 4 | 450 (x6), 300 (x8), 298, 299, 299, 443, 444, 293 | $P < 10^{-4}$ (x15), $P < 10^{-2}$, (x3), .0126, .4757 | 1.54 |

**Figure 3.**
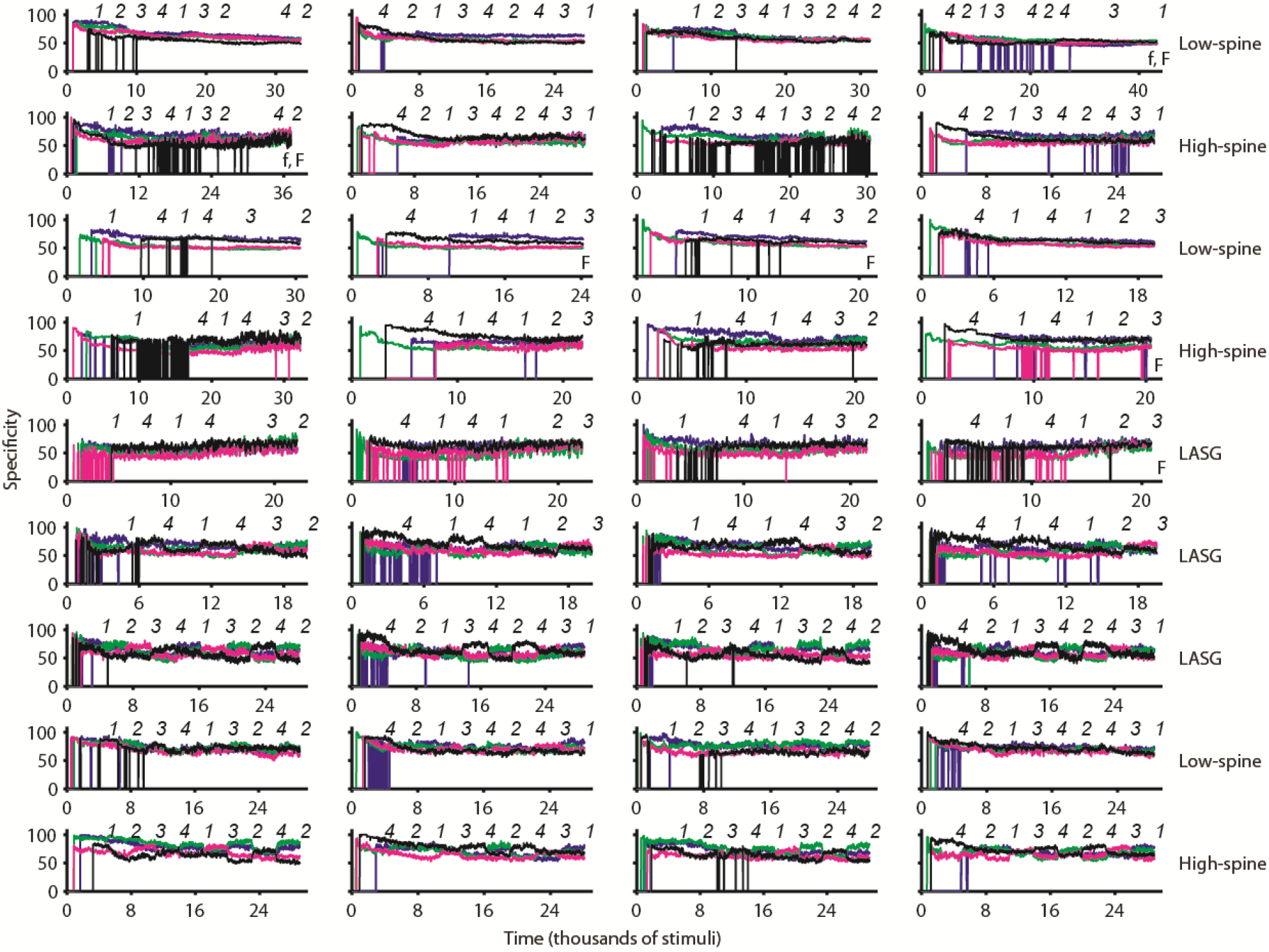
Response specificity measurements, for all simulations that exhibited successful learning. Plot elements match those of Figure 1C. The first (last) two columns of plots correspond to ISN1 (ISN2). The plots in the first five (last four) rows correspond to simulations without (with) PING. For the plots in each row, the candidate synapse generation regime is as indicated at right. “F” denotes simulations for which there was an instance of total learning failure prior to the depicted result, while “f” denotes simulations that contain a partial learning failure (defined as exhibiting exceptionally weak response specificity towards certain configurations, even after training).

**Figure 4.**
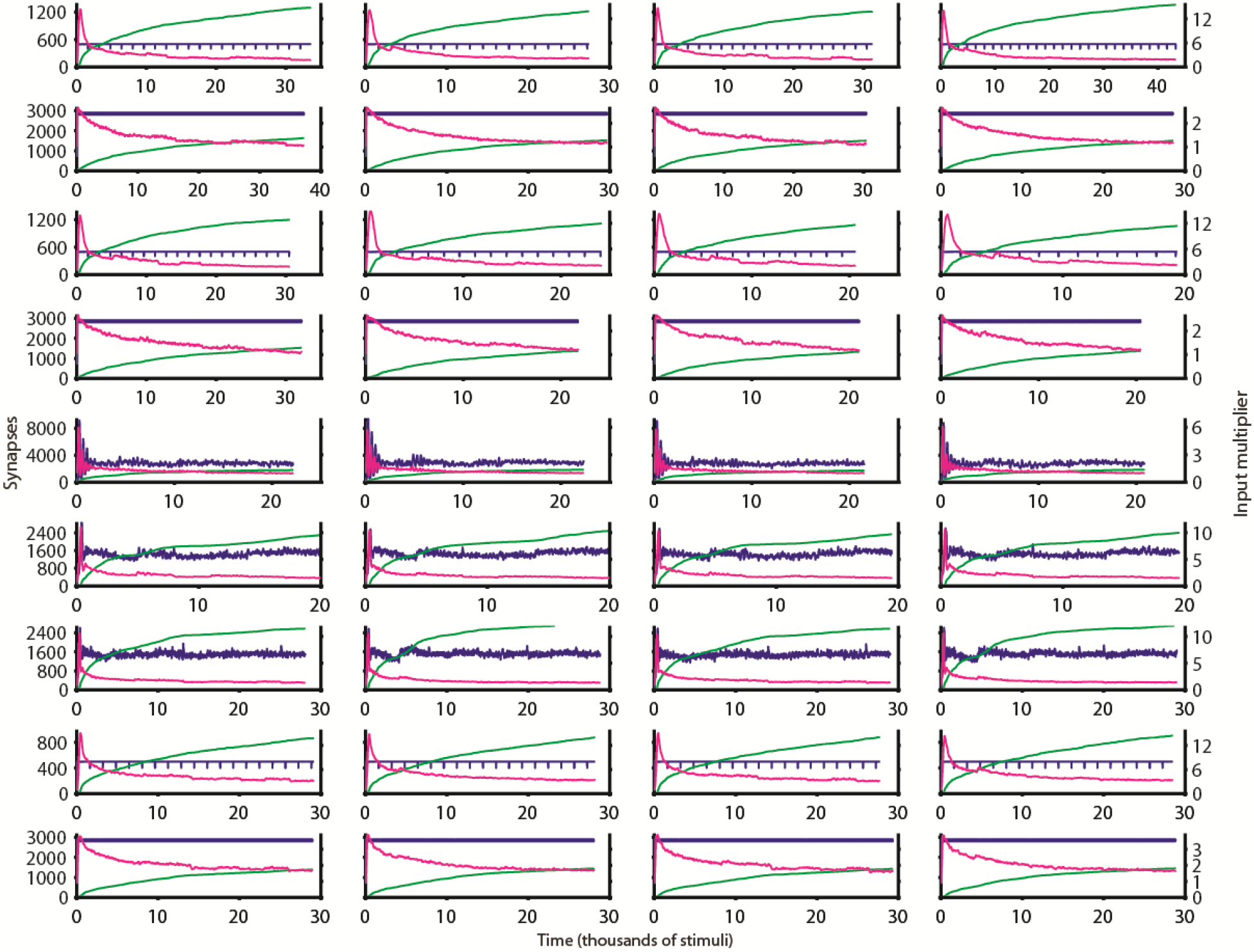
Time course of connectivity development for all simulations that exhibited successful learning. Data in each plot derive from the same simulation as the equivalently-positioned plot in Figure 3. Subplot elements match those of Figure 1B.

### Learning-accelerated spine generation overcomes anterograde interference, sacrifices response specificity

An important consideration that shapes our examination of LASG’s effects is that LASG removal markedly reduces candidate spine generation. Accordingly, all no-LASG simulations are carried out in both a low-candidate regime (default baseline candidate generation rate) and a high-candidate regime (elevated baseline generation rate that produces candidate synapse populations size-matched with the LASG regime). This informs our understanding of which effects are specifically attributable to LASG’s absence rather than to anemic candidate generation.

The major outcome of our comparisons of the LASG, high-candidate, and low-candidate regimes is that candidate synapses in general and their LASG-driven generation in particular are detrimental to response specificity. This is the result that we would expect, upon considering biologically-observed pruning and previous theoretical work that demonstrates pruning’s noise-reduction(29, 30) benefits. Specifically, in PING’s presence, for all four input orderings, applied to both ISNs, response specificity is higher for both no-LASG conditions than for the LASG condition (Table 2, rows 3 and 4). Additionally, in PING’s absence, this holds true for the 6-epoch input orderings (Table 2, rows 6 and 7), and also arguably holds true for the 9-epoch input orderings, for which learning fails altogether during the LASG regime. Meanwhile, there is some evidence for a response specificity difference between the low- and high-candidate regimes that favors the former (Table 2, rows 5, 8, and 9; unpublished raw data). Finally, we find that response specificity is very often improved by removal of candidate synapses (Table 2, rows 10 and 11); the simulations that show the least improvement all correspond to the low-spine condition and can therefore be regarded as the exceptions that prove the rule.

LASG clearly comes with a price, and one may wonder what benefit, if any, it provides that can be replicated *in silico*. Though we observe no benefit in simulations without PING, it is clear by inspection (Figure 5A) that of the three PING simulations shown, the LASG regime exhibits the strongest configuration-specific response specificity shifts. To study these shifts, we proceed as follows. First, we define the “response competitiveness” (RC) as the difference between the response specificity to an epoch’s assigned configuration and the average response specificity to the other configurations. Second, because anterograde interference favors a simulation’s first-experienced input configuration, we focus our analyses on the first two epochs in which each of the three non-initial configurations are experienced. Third, because RC appears to increase over the course of an epoch, we split each epoch into segments, and within each of these windows average the measured RC values. Fourth, because the process of RC development appears to repeat itself for each of the non-initial configurations, we opt to examine RC within each epoch segment by pooling across the three non-initial configurations. Fifth, RC values corresponding to epoch segments of the same input configuration ordering, compared across spine generation regimes, can be regarded as matched pairs.

**Figure 5.**
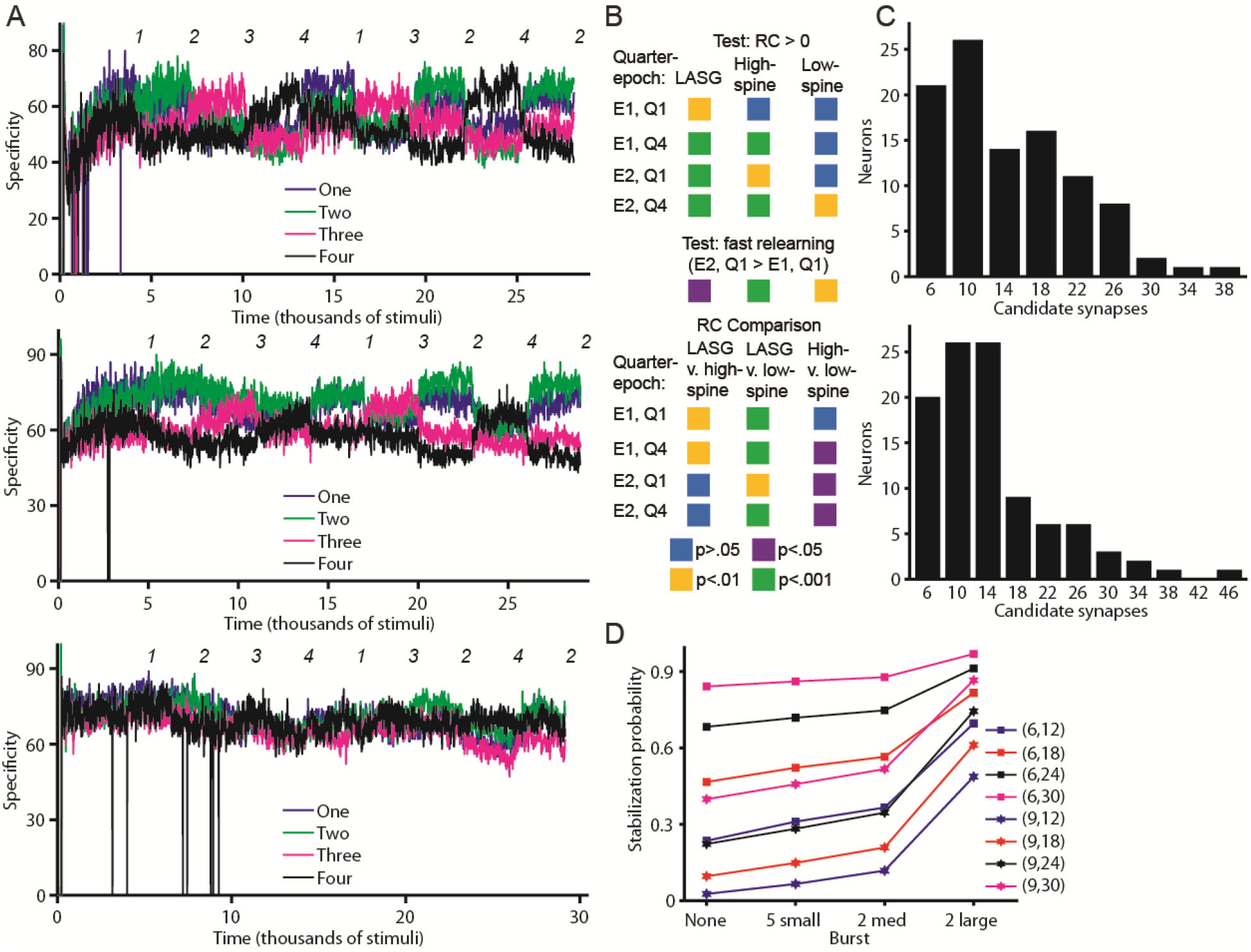
LASG ameliorates anterograde interference (in networks with PING). **A** depicts the typical tuning specificity behaviors exhibited by networks simulated with the LASG (top), high-spine (middle), and low-spine (bottom) candidate synapse generation regimes. The depicted simulations were run with ISN1, and the depicted specificities were measured with candidate synapses intact; otherwise, elements of each subplot match those of Figure 1C. **B** depicts the results of response competitiveness analyses across the three candidate generation regimes; specifically, it depicts the statistical significance of one-sample and matched-pair t-tests (df=11) conducted using vectors of RC values. **C** depicts typical histograms of candidate synapse counts across the population of output neurons. For the values *v* denoted on the x-axis, histogram bins correspond to the interval [*v*-1,*v*+2]. The two network state snapshots depicted respectively represent the midpoint and endpoint of the 5^th^ training epoch of the simulation whose tuning specificity is depicted in panel A (top). **D** depicts the probability that an input will be learned during a time interval, with that probability depending on the number of input-specific candidate synapses whose concurrent presence is needed for learning (N), the time window’s average number of candidate synapses (C), and the clustering of candidate synapses into bursts (x-axis). The lines spanning the x-axis are labeled according to (N, C).

Our RC data, distilled as described, reveal that RC emerges most quickly in simulations with LASG, that high-spine simulations eventually catch up, and that in low-spine simulations RC is weak and slow to develop but develops eventually. An additional discovery enabled by RC analysis, congruent with biologically-observed slow initial learning and faster relearning of sensory module mappings, is that RC emerges more quickly during a network’s second encounter with an input configuration than during the first. All of these conclusions are supported by multiple segmentations of the epochs, but for simplicity, we demonstrate them here by segmenting epochs into quarters and using each epoch’s first and fourth. Outcomes of individual analyses performed are depicted in Figure 5B.

It is somewhat surprising that the LASG regime exhibits faster RC development than the high-candidate regime despite the latter producing a greater overall number of candidate synapses; this result can, however, be explained by the LASG regime’s candidate synapse bursts. Specifically, while the distribution across neurons of candidate synapses is uniform in LASG’s absence, LASG will occasionally drive select neurons to produce many candidates (Figure 5C). We presume that, because of the model’s moderately long synapse lifetime, the epoch’s most frequently recurring input correlations will probably overlap with the bursts, and thus the synapses that transmit those inputs will be stabilized. We probe this intuition by computing (details in *Methods*) the effect of candidate synapse bursts on the probability that a specific cluster of synapses projecting to a neuron will be stabilized within a given time window. Consistent with LASG’s hypothesized effect, we find (Figure 5D) that bursts greatly increase learning probability, even when the candidate count is low overall.

### Previously-reinforced memories produce anterograde interference via Hebbian competition

To better understand our system’s apparent exhibitions of anterograde interference, we create a simplified network in which 100 linearly-arranged input neurons compete for connectivity to a single output neuron. More specifically, two clusters of neurons with correlated activation are defined as memories, while the other neurons exhibit individual activation independently of each other or of the clusters. This setup allows us to examine synaptic weight changes under each of the following three memory competition conditions: equally sized clusters and weakly-initialized synaptic weights, equally sized clusters with the first cluster’s weights initialized as being fully reinforced, and an enlarged second cluster with the first cluster’s weights initialized as being fully reinforced. Simulation details are provided in *Methods*. Upon running 20 short simulations under each condition, we find the second cluster’s simulation-end weights to be particularly revelatory. Specifically, their average weight across all simulations remains close to the weak initial weight in experiment condition two, is approximately twice (thrice) as high in experiment condition one (three), and the rank sum test (n=20) confirms that the weights (averaged within a simulation) produced by experiment condition one differ significantly from both condition two’s weaker weights (p=.0329) and condition three’s stronger weights (p=.0364). These results corroborate our notions that strong synapses from a simulation’s previous epochs generally interfere with the present epoch’s learning, and that large groups of synapses (e.g. those generated by LASG) with correlated signaling are able to overcome that interference.

### Individual synapse association with a specific input configuration manifests in weight fluctuations and across multiple simulations

From *in vivo* observations of correspondence between memories and specific synapses (31), and our *in silico* observation of epoch transitions being accompanied by accelerated synapse stabilization and of (in simulations with PING) the correlated fluctuations of configuration-associated synapse groups, we infer that the existence of and heaviest signaling through certain synapses is driven by a specific input configuration. To corroborate this inference at the level of individual synapses, we analyze (see *Methods* for details) the data from the first four epochs of the simulations depicted in Figure 3’s 5^th^ row. This analysis helpfully restricts us to the synapse sets of just two input configurations 1 and 4, which are conveniently the configuration pair whose stimuli differ the most from each other. We find that for both configurations, the corresponding sets of synapses overlap strongly across the two simulation runs with the same ISN; meanwhile, there is no statistically significant strong overlap between synapse sets corresponding to different configurations (Table 3). This finding demonstrates that the stabilization of at least some synapses is preferentially associated with a specific input configuration. Next, our notion that an input configuration’s reinforcement and suppression effects are synapse-specific is bolstered by an examination of within-epoch synaptic weight fluctuations, which reveals that individual synapses tend to remain consistently strong or consistently weak (Figure 6A). Moreover, because synapses are clearly classifiable as strong or weak within an epoch, we are able to distill eight groups of synapses (one group for each combination of input ordering, input configuration, and ISN) that are strong specifically during epochs with their preferred input configuration, and weak otherwise. All groups of synapses exhibiting this weight shift pattern are much larger than would be expected (Table 4) if each epoch’s strong-or-weak synapse assignments were made at random.

**Table 3.** Analysis of overlap between configuration-specific synapse sets drawn from different simulations

| ISN | n | Set 1 | Set 2 | E | A | p |
| --- | --- | --- | --- | --- | --- | --- |
| 1 | 11235 | 1, 1414 | 1, 4141 | 59 | 111 | < .001 |
| 1 | 11235 | 4, 1414 | 4, 4141 | 46 | 89 | < .001 |
| 2 | 11174 | 1, 1414 | 1, 4141 | 55 | 122 | < .001 |
| 2 | 11174 | 4, 1414 | 4, 4141 | 57 | 117 | < .001 |
| 1 | 11235 | 1, 1414 | 4, 4141 | 174 | 149 | .984 |
| 1 | 11235 | 1, 4141 | 4, 1414 | 16 | 17 | .299 |
| 2 | 11174 | 1, 1414 | 4, 4141 | 173 | 168 | .655 |
| 2 | 11174 | 1, 4141 | 4, 1414 | 18 | 14 | .810 |

The ordered pairs in columns "Set 1" and "Set 2" correspond to (configuration, input ordering), and denote the two synapse populations being compared. The values of n and E, and A respectively correspond to the total number of axons, the expected number of synapses common to the synapse populations (assuming that the membership of each is selected at random from the axons), and the actual number of common synapses. P values correspond to the one-sided hypergeometric test.

**Table 4.** Existence of well-defined synapse populations strengthened specifically in the presence of their corresponding input configuration

| ISN | Configuration | Ordering | n | E | A | p |
| --- | --- | --- | --- | --- | --- | --- |
| 1 | 1 | 1414 | 1402 | 49 | 95 | < .001 |
| 1 | 1 | 4141 | 470 | 84 | 98 | < .001 |
| 2 | 1 | 1414 | 1391 | 39 | 91 | < .001 |
| 2 | 1 | 4141 | 443 | 54 | 71 | < .001 |
| 1 | 4 | 4141 | 1393 | 40 | 74 | < .001 |
| 1 | 4 | 1414 | 373 | 62 | 75 | < .001 |
| 2 | 4 | 4141 | 1391 | 38 | 92 | < .001 |
| 2 | 4 | 1414 | 457 | 69 | 80 | < .001 |

Each row corresponds to a specific simulation's configuration-specific synapse population; column n states its size. As for the population of synapses whose epoch-specific strength tracks the configuration's presence, E and A respectively correspond to its expected (assuming random strong-weak synapse assignments) and actual size. P values correspond to the one-sided binomial test.

**Figure 6.**
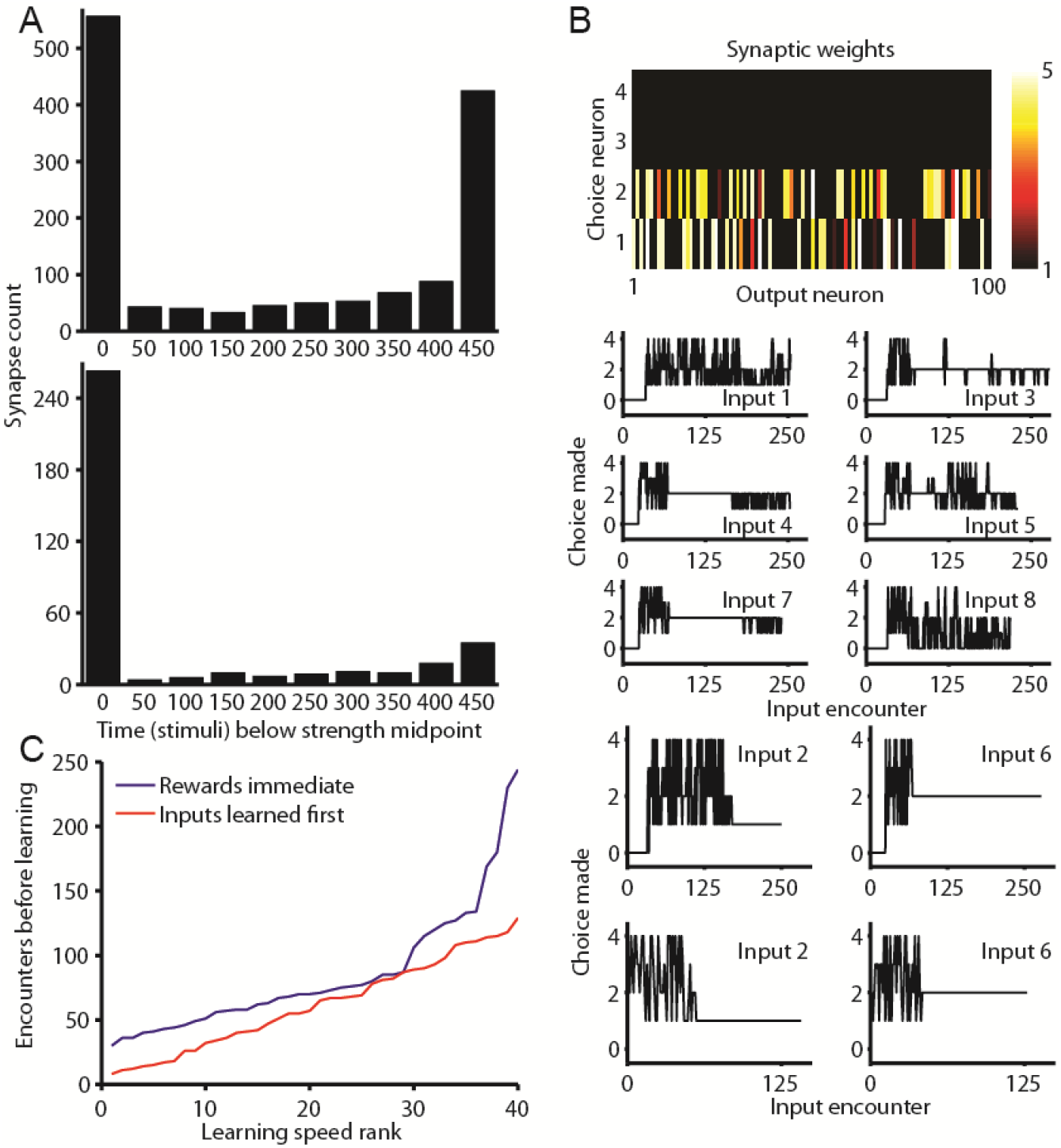
Analysis of synapse cluster behavior: correlated weight shifts and enablement of stimulus-specific reinforcement learning. **A** depicts the distribution across synapses of time spent below the midpoint of the synapse strength range. Data are drawn from the window corresponding to the presentation of the last 500 stimuli of epoch 4 (ISN1, input ordering 1414). Top and bottom subplots respectively correspond to synapses stabilized during training with input configurations 1 and 4. **B** depicts the network’s learning to associate individual stimulus bars with choice options. The color map (top) depicts a representative simulation’s end-state synaptic weights between the output and choice neurons. For this same simulation, the six plots in the panel’s middle segment indicate the choice neuron activated during encounters with inputs not reward-associated with a specific choice. Meanwhile, the network’s choices during encounters with the reward-associated inputs are depicted in the bottom segment’s top two plots. The bottom two plots show the same, but for a simulation in which choices were presented at the midpoint rather than from the outset. **C** depicts, for simulations in which choices are presented either immediately or at the midpoint, the number of input encounters necessary before the input is consistently associated with the rewarded choice option. For both choice presentation conditions, the data shown are pooled from both ISNs (10 simulations for each condition-ISN combination) and both reward-associated inputs in each simulation. Learning requires significantly fewer reward-informed input encounters when choices are presented at the midpoint vs. at the outset (two-sided rank sum test, p=.0413, n=40).

### Synapse groups that represent input correlations enable performance of discrete memory tasks

Finally, the emergence of Hebbian ensembles is a memory acquisition concept whose relevance extends beyond questions of sensory map formation. As a first exploration of the applicability of our structural plasticity approach to discrete learning tasks, we simplify our simulation so as to learn just eight bar inputs (Figure 1A, Test stimuli, input configuration 1), add a set of “choice” neurons to our network, and reward the association of input 2 with choice 1 and of input 6 with choice 2. (Network and learning mechanism details are provided in *Methods*.) Stimulated in this manner, the network develops reinforced connections from mostly non-overlapping sets of output neurons to the 1^st^ and 2^nd^ choice neurons (Figure 6B, top), learns to consistently (and, respectively) associate inputs 2 and 6 with choices 1 and 2 (Figure 6B, bottom), and develops a bias towards choices 1 or choice 2 when presented with any of the other inputs (Figure 6B, middle). These results demonstrate that neuron clusters corresponding to repeatedly-encountered discrete entities (in this case, bar stimuli) are able to operate as a group (in this case during an associative learning scenario). Furthermore, the network’s preference for choices 1 and 2 in response to the six no-reward inputs can be regarded as an exhibition of capacity to generalize based on other reward-associated experiences. We next demonstrate that the learning of associations between rewarded choice options and discrete structured inputs is necessarily preceded by the development of representations of the discrete structured inputs (i.e. the bars). With both ISNs, we run simulations that vary according to whether choice-making and reward information availability are present from the beginning of the simulation (Figure 6B, bottom, top two plots), or introduced at the midpoint (Figure 6B, bottom, bottom two plots). What we find is that in the former simulations, choice behavior is absent (i.e. choice 0) early on, presumably while the output module clusters corresponding to the bars are being formed, but that in the latter simulations, the number of reward-informed input encounters needed for the preferred choice associations to be learned is much lower (Figure 4C). Intriguingly, our network’s ability to encode environmental correlations and subsequently ascribe them reward valence is suggestive of latent learning; we believe that a somewhat more elaborate network, endowed with our structural plasticity modeling scheme, would be able to exhibit even richer cognitive bootstrapping capacity.

## Discussion

In this paper, we address the difficult problem of learning and selectively accessing multiple overlapping memory representations by putting forth a solution that incorporates biologically-inspired mechanisms, those of structural plasticity and inhibition-mediated input filtering, not typically incorporated into synthetic memory systems. This exercise suggests memory system design strategies and reveals design trade-offs that inform both the engineering of synthetic systems and our understanding of biological ones. It also presents an approach to structural plasticity modeling with clear potential to open up several avenues of neuroscientific inquiry.

Our system addresses several information processing challenges faced by learning systems. A first is the challenge of long-term retention (3), which in some contexts manifests as retrograde interference (6). Our biomimetic strategy for permanent storage is permanent stabilization of the pertinent synapses; accordingly, in both our network and biologically-embedded ones, memory acquisition correlates with accretion of permanent synapses. Our confidence in this strategy is reinforced by the contrast between models that employ slowly-changing synapses (5, 6, 32) and models whose synapses have state descriptor variables that explicitly tend to stay constant after initial learning (33, 34). A second challenge concerns a different type of interference, anterograde, in which a network’s previously-acquired memories hinder the subsequent acquisition of memories that a tabula rasa network would have no trouble acquiring. Notably, in Knoblaugh’s structural plasticity model (29), anterograde interference arises from reduced availability of memory-unassigned synapses, while in ours, the signaling through the synapses of previous epochs creates broad activation that degrades response specificity. Fortunately, our model’s form of anterograde interference is amenable to being overcome by PING, which restores stimulus response sharpness, and by LASG, which recruits entire ensembles of synapses to transmit the desired signal. A third challenge, also overcome by PING and also an issue of memory interference, is recall differentiation between memories that are very similar, possibly to the point of overlap. Over the course of an epoch, PING weakens the synapses corresponding to the interfering representations, and the network’s resultant signaling is specifically favorable to each epoch’s assigned input configuration. Mechanisms that allow for engram-specific reinforcement, even when their representations overlap, are at work in biological systems as well (31). Finally, a fourth design challenge is learning flexibility; our system’s flexibility derives from the random connectivity of the ISNs, which enables the capture, by some subset of potential synapses, of whatever correlations present themselves in the environment. We hypothesize that such wiring schemes are implemented in association cortices and other subnetworks in which there is a requirement for breadth of learnable input correlations. Notably, spatial randomness of projections that allows for the self-organized development of input-representation correspondence is a design feature of the piriform cortex (35). Similar randomness is found in the organization of orientation-tuned neurons in mouse visual cortex; in animals with larger visual cortex, more elaborate organization schemes are thought to be driven by minimization of biological wiring length rather than by information processing considerations (36).

Some of the challenges faced by learning systems arise from inherent tension between learning objectives; several of our system’s learning outcomes reveal these tensions, and several system features are in turn revealed to play a role in mitigating them. For example, we see in Figure 3A (top plot in particular) that emphasizing the signal of the current epoch’s configuration comes at the cost of representation specificity for the other configurations, the recovery of which requires a non-zero amount of relearning time. The weakening of synapses corresponding to off-epoch configurations is mitigated by their long-term stability, which allows them to be reinforced in parallel when the need arises. The difference between serial (parallel) reinforcement during (re)learning is also revealed by our connectivity development plots (Figures 1B, 2C, green traces). Notably, our model’s learning kinetics are consistent with findings that human subjects who experience a visual input distortion to which they had previously adapted are able to re-adapt more quickly than initially, but do not re-adapt instantaneously. The other design tension we encounter arises between learning flexibility and the desirability of pruning candidate synapses for the sake of noise reduction. The noise reduction theme arises quite consistently across computational studies of structural plasticity: we found that candidate synapses are a noise source (Table 2, rows 10 and 11), Spiess et al (30) found that well-pruned networks are less noisy and therefore learn faster than their unpruned counterparts, and Knoblaugh et al (29) found that networks have greater information capacity per synapse when some synapses can be pruned altogether rather than simply weakened. In our system, the allowance of a limited population of weak exploratory candidates is sufficient to allow for learning flexibility, but not enough to enable rapid learning. In both our model and Knoblaugh et al’s (29), candidate synapses generation is a learning speed bottleneck, because the synapses whose stabilization would constitute learning are not all simultaneously present, and learning must therefore take place over multiple exposures to an input. Our characterization of the “spacing effect” as the outcome of a candidate synapse availability bottleneck is intriguingly complementary to explanations that emphasize memory reconsolidation dynamics (37) or short-term desensitization of learning-correlated intracellular signaling (38). LASG, of course, is our system’s mechanism for overcoming the learning speed price of noise reduction. It is a particularly clever mechanism because it allows the system to temporally separate the pursuit of pruning and the pursuit of learning; in fact, because of the spatiotemporal specificity of LASG, the memories blurred by bursts of spine generation are precisely the memories whose representation fidelity doesn’t matter at the time of the burst.

Our final comment on the engineering implications of our work is to highlight that our results, together with the above catalog of our system’s computational feats, vividly demonstrate that many mechanisms (PING, LASG, pruning, Hebbian competition, etc.) work in concert to perform the many operations that cognitive processing requires.

As a neuroscience tool, our structural plasticity modeling method has value that Feynman (39) would appreciate: it is uniquely useful for replicating certain phenomena *in silico*. One such set of phenomena is the circadian features of synaptic turnover, e.g. long candidate synapse lifetimes and alternating periods of activity and synapse accumulation vs. quiescence and pruning (40). A second set of phenomena our method can (likely) replicate are the various manifestations of cognitive bootstrapping and category learning. A third candidate domain for our method’s application is the study of graph structure development and the emergence of constructs such as hub neurons (41). Modeling of any of the above processes would benefit from learning task and network architecture design being guided by, and then ground-truthed against, real-world learning outcomes; in the present study, the guiding real-world outcome was slow learning and fast relearning of sensory module mappings.

Perhaps the most fruitful future application of the tools and paradigms presented here will be to the interpretation and conceptual organization of experimental neurobiology findings. One of two reasons why is that in modeling studies, all aspects of system behavior are measurable, which in turn allows for the precise definition of information processing metrics (e.g. response specificity, input encounters required for learning). Secondly, because of this measurability, simulations built to achieve a specific cognitive task offer stages on which mechanisms can quantifiably showcase their contribution to task-relevant information processing. It is with these opportunities in mind that we consider how distinct memory modification events (e.g. acquisition, reconsolidation, and extinction) are correlated with distinct ensembles of intracellular signaling and gene expression responses (42–44) and are distinctly affected by Arc mutations (45). Similarly, events that trigger different durations of neural activity are processed differently; they trigger the expression of different sets of signals and genes (46). Our own study’s rules for cell and synapse modification could certainly be expanded to incorporate what is known about the above biological scenario differences, thereby allowing the biology to be understood in the languages of filtering, of representation fidelity, of environmental estimation, and of other emergent information processing outcomes. Tolman (47) wrote in 1938 that theories complement catalogs of experimental permutation results by serving as concise descriptions of the world with great predictive generality; a computational account of the extant catalog of learning-associated neurobiology would provide precisely these benefits.

## Acknowledgments

This work was funded by National Institutes of Health grant 5T32HD060555 and by the University of California, Irvine Eugene Cota-Robles Fellowship.

## Methods

### Network

To model the development of mappings between multiple input modules and an output module to which they commonly project, we use the network sketched in Figure 1A (Network). This network is composed of integrate-and-fire neurons, arranged in three grids of the indicated dimensions. Input neurons extend “axons” (blue arrows) to output neurons; input-output neuron pairings are assigned an axon with 20% probability. These are the network’s only connections. At the beginning of every simulation run, they are instantiated as “potential” synapses (defined below). In this study, we designate an ensemble of axons as an “initial state network” (ISN) for an, and refer to the two we generate as ISN1 and ISN2.

### Stimuli

The network is trained with stimuli whose basic form is three randomly-placed, identically-oriented, non-overlapping bars (Figure 1, Sample input). The size (in neurons) of individual bars is 1×5; that is, bars are implemented as current injection (details below) to five linearly adjacent neurons. New stimuli are generated on an ongoing basis throughout a simulation, and a transformed version thereof is presented to each of the input modules. The four transformations used in our study are shown in Figure 1A (Input configurations): both modules experience identically-positioned 1) vertical or 4) horizontal bars, or else module 1 experiences vertical bars and module 2’s input is module 1’s but either 2) mirrored across the middle column of neurons or 3) rotated counterclockwise 90 degrees.

### Response specificity

At regular intervals, the network’s response specificity to different inputs is measured. Specifically, for every one of the input configurations used in our study, eight single-bar stimuli (Figure 1A, Test inputs) are sequentially presented. Assigning to the input grid’s bottom-left neuron the coordinates (1, 1), the coordinates of the bottom neuron of the baseline (i.e. prior to any configuration-specific transformation) 1×5 vertical bar stimuli are (3, 2), (3, 10), (6, 2), (6, 10), (10, 2), (10, 10), (13, 2), and (13, 10). Then, for each configuration’s stimuli octet, output module response specificity is quantified as follows. First, output neurons are grouped according to which of the eight stimuli elicited from them the most firing (in case of ties, neurons are assigned to multiple groups). Response specificity is deemed absent, if for any of the stimuli there are no output neurons that exhibit a preferential response. Usually, however, the next step is to pool together the firing of neurons in each group *s*, and compute the total spikes elicited by all the stimuli (*T*_*s*_) and by the preferred stimulus (*P*_*s*_). Whole-network response specificity is thus defined as 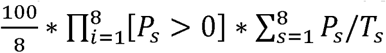.

### Time

The basic unit of time in our study is the period during which a specific stimulus is presented. (Thus, for example, simulation durations are reported in stimuli.) In practical implementation, this time unit is in turn further discretized. During the first 90% of a stimulus period, the stimulated input module neurons are injected with current; this is followed by a brief moment of input silence.

### Neurons

The voltages of our model’s input neurons update according to Eq. 1 (in which *i, t, τ*_*V*_, and *S* respectively represent the neuron’s index, the time, the voltage decay rate, and, when applicable, the bar stimulus), except when 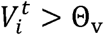 (Θ_v_ is a firing threshold), in which case 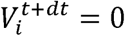. Similarly, output neuron voltages, below threshold, are updated according to Eq. 2 (in which *j, N*_*pre*_, 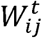, and [] respectively correspond to the neuron’s index, the number of input neurons from which neuron j receives projections, the synaptic weight between neurons *i* and *j*, and a truth operator). In Eq. 2, synaptic inputs are scaled by a homeostatic input multiplier, *F*, that itself updates according to Eq. 3 and Eq. 4 (in which *r*_*j*_, *r*_*r*_, *r*_*o*_, and *τ*_*F*_ respectively represent a measure of recent activity, its decay rate, its set point, and the rate constant of input multiplier change).

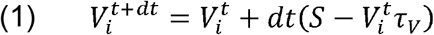

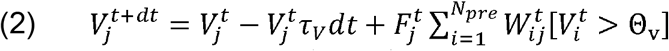

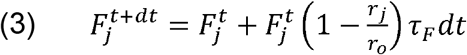

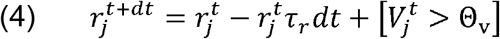

### Synapses

A synapse is described by its Hebbian reinforcement 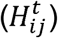, activity-dependent depression 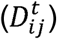, weight 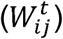, and state 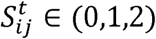. Hebbian reinforcement’s magnitude is an exponential function of the firing time difference, truncated at *q*, between the pre- and post-synaptic neurons (Eq. 5; *t*_*i*_ and *t*_*j*_ are the most recent firing times of these neurons). Meanwhile, activity-dependent depression, which takes the form of a weight decrement L, occurs if presynaptic firing is not followed by postsynaptic response within time 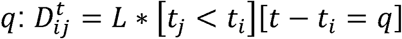. Combining the above, weights are updated according to 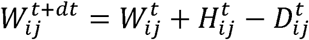, subject to 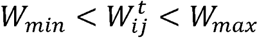 (*W*_*min*_ and *W*_*max*_ are weight bounds).

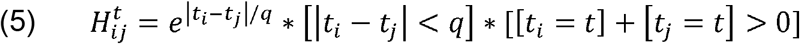

Every output neuron has a candidate integrator 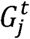, which updates according to Eq. 6 (in which *G*_*r*_ represents the strength of LASG), unless 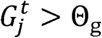 (the synapse generation threshold), in which case 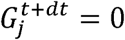 and a candidate is generated from a randomly selected potential synapse.

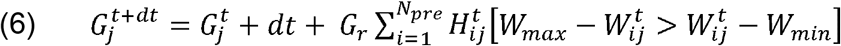

The three possible values of *S* denote synapses that are potential (0), candidate (1), or long-term stabilized (2). Transitions amongst states are as indicated in Figure 1A (Synaptic states). Candidate synapses (*S* = 1) are additionally described by their total experienced Hebbian reinforcement 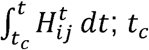 is the candidate’s time of creation) and a collapse countdown timer *µ* with an initial value *µ*^*I*^. Naturally, 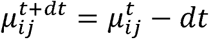. Candidates transition to either the potential (if 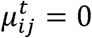) or stabilized (if 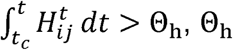 is a stabilization threshold) state, depending on which condition is met first.

### Structure of simulations

Each of our simulations is composed of epochs, and epochs in turn are defined as sequences of equivalently-configured stimuli. We structure our experiments around configurationally-homogeneous epochs because our aim, after all, is to model the network’s learning to differentiate between stimuli in each of the different configurations.

Within an epoch, system state measurements are recorded after every 10^th^ stimulus: total candidate and stabilized synapses, synaptic weights, average input multiplier, and the network’s response specificity to the test inputs in each of the four input configurations. Response specificity measurement proceeds as follows: a copy of the network is made in which connectivity is frozen, all neuron voltages are set to resting potential prior to the presentation of each test stimulus, and current is injected during the entire stimulus window with no input silence at the end.

The details of every simulation are determined according to the specific experimental objective; generally, discovering how certain modifications, e.g. to parameters, affect learning. With this objective in mind, all of our simulations are done in quadruplicate across two ISNs and a pair of either short or long input orderings (a simulation’s input ordering is defined as the number of epochs and the input configuration assigned to each), with the ISNs and input orderings in turn being reused across experiments (specific input orderings used are those indicated in Figure 3). This regime reveals what learning changes generalize across ISNs and input orderings, while removing these variables as potential confounding factors in result interpretation.

Whereas each epoch is dedicated to the network’s acquisition of response specificity to stimuli in a certain configuration, and whereas learning typically drives the stabilization of new synapses, we regard learning as complete when three conditions are met: that the number of stabilized synapses during the most recent 250 stimuli comprises <2% of the total stabilized, that over the previous 100 response specificity measurements (for the configuration corresponding to the present epoch) no more than 5 return a score below 50, and that the epoch’s run time is at least 2000 stimuli. Response specificity measurements are made with all candidate synapses removed, so as to confirm the presence of a set of stabilized synapses that effectively differentiate between the Test stimuli. If the conditions are satisfied, the epoch runs for another 1000 stimuli, and then the simulation transitions to the next epoch. Alternatively, if stabilization is not achieved after 8000 stimuli, the epoch proceeds for another 1000 and is then terminated. At this point, if the latest epoch exhibits weak but intact response specificity, the epoch is regarded as a partial learning failure and the simulation transitions to the next epoch. If response specificity is frequently absent, however, we regard the epoch as a total learning failure, and re-run the planned simulation from the beginning.

### Pyramidal-inhibitory network gamma (PING)

We implement PING by adding to the above network an inhibitory cell that is reciprocally connected to all of the output module’s excitatory neurons. Rather than being subject to structural and strength plasticity, these connections are permanently existent and statically weighted (*W*_*OP*_). The voltage of the PING-generating neuron (*P*), below threshold, is updated according to Eq. 7, and output neurons are now updated according to Eq. 8.

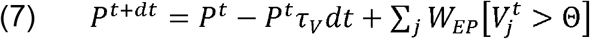

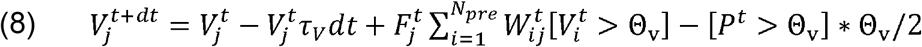

### Response specificity comparisons

We accomplish the comparison of response specificities across different experimental conditions using the following procedure. First, unlike the response specificity measurements made with candidate synapses removed, as described above, our comparisons between different candidate generation regimes are based on measurements made with candidate synapses intact. Second, because response specificity changes over the course of the typical epoch, because the network’s state is relatively stabilized at an epoch’s end, and because epochs vary in duration, we choose to standardize comparisons between simulations by focusing on each epoch’s final 50 response specificity measurements. Third, because each epoch’s training is intended to develop response specificity to stimuli in a particular configuration, we focus our analysis on the measured specificity values of each epoch’s assigned configuration. Fourth, because our objective is to compare response specificities across different experimental conditions, we pool each simulation’s epoch-end specificity values into a single vector. These vectors allow for simulations that differ in one specific way, but that share ISNs and input orderings, to be directly compared; notably, equivalently-positioned values in each vector will have been generated during equivalent epochs! The fifth step of our process, then, is to clean the vectors of measurements of zero-value specificity instances, which would otherwise confound comparisons; we delete the equivalently-positioned measurement in both vectors being compared, so that they may maintain the same proportion of measurements derived from each input configuration. Finally, even after cleaning, many vectors of response specificity values, as well as element-by-element differences between pairs thereof, returned Liliefor’s test p-values below. 05; so, for consistency, all response specificity comparisons were performed using the rank sum test (as indicated in Table 2).

### Synapse cluster stabilization probability computation

To examine the effect of candidate synapse bursts on the probability that a specific cluster of synapses, transmitting a common stimulus and projecting to the same neuron, will be stabilized within a time window, we proceed as follows. We divide the time window into 20 intervals and distribute and distribute a fixed number of candidate synapses across the intervals. With no bursts, candidates are distributed uniformly. Meanwhile, the five small (two medium, two large) bursts condition shifts to the burst intervals 4 (3, 6) candidates from each of the remaining intervals. Then, for each interval, the probability that the candidate synapse set, drawn from a total pool of 100 potential synapses, will encompass the synapse cluster of interest (there is an assumption that stabilization of the cluster requires that all of its members be coincidentally present) is calculated using the hypergeometric distribution. Labeling each interval’s stabilization probability as *P*_*i*_, the stabilization probability of the time window as a whole is 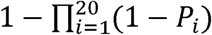.

### Single-target Hebbian competition

Here, our aim is to demonstrate, at the level of a single cell, that previously-acquired memories (i.e. clusters of reinforced synapses that exhibit correlated firing) interfere with the acquisition of new memories (i.e. joint reinforcement of other synapse clusters), but that this interference can be overcome if the group of synapses representing the previously-unreinforced memory is large. To conduct this inquiry, we first simplify the baseline (i.e. the model as described prior to the introduction of PING) modeling approach by reducing it to one output neuron and a single row of 100 input neurons, and dispensing with the synapse generation process in favor of simply creating a stabilized synapse from each input neuron to the output neuron. In this simplified model, we approximate memories as clusters of synapses that always fire jointly. We also revise the process of selecting which neurons are stimulated during each stimulus period: now, every memory cluster, as well as every unaffiliated neuron, is stimulated with probability *P*(*active*). Finally, we adjust *r*_*o*_ (output neuron’s homeostatic activity set point) downward, to induce Hebbian competition even though the output neuron is receiving less input than in the baseline model.

Having developed a model tailored to the study of interaction amongst clusters, we define three experimental conditions and under each condition, run the model for 20 trials of 400 stimulus periods each, and record each trial’s end-state synaptic weights. The first cluster’s size is always 5; the second cluster’s size is 5 in conditions 1 and 2 and 7 in condition 3. Meanwhile, synaptic weights are generally initialized at *W*_*min*_, but in conditions 2 and 3 the first cluster’s weights are initialized at *W*_*max*_.

### Analysis of configuration-specific stabilization and weight fluctuation

Our goal is to identify synapses that correspond to a particular configuration, and study their weight shifts across multiple episodes. Because most configuration-specific synapses are stabilized during the network’s initial encounter with a particular configuration, and because interpretation of weight fluctuations for synapses stabilized later is not straightforward, we filter each simulation’s stabilized synapse set so as to isolate for each configuration the synapses stabilized during the initial encounter. Next, we extract the strengths of these synapses from the time window corresponding to each epoch’s final 500 stimuli, because at this point network structure and synaptic weights have presumably annealed. Within this time window, synapses that are above or below the strength range midpoint for over 90% of the duration are respectively classified as strong or weak.

Consider the input ordering XYXY. Let us denote configuration X’s synapses that are strong during epoch one as *X*_1,*S*_, that are weak during epoch two as *X*_2,*W*_, and proceed similarly for the remaining epochs and for the synapses of configuration Y (note: *Y*_1,*S*_ and *Y*_1,*W*_ are both the empty set). Further, let the total populations of the two configurations be denoted as *X*_*T*_ and *Y*_*T*_. Finally, let us denote the size of each of these populations via the prefix *s*, e.g. *sX*_1,*S*_. With this notation established, we see that the sets of population X and Y synapses that exhibit epoch-specific strength fluctuations can be respectively denoted as *F*_*X*_ = *X*_1,*S*_ ∩ *X*_2,*W*_ ∩ *X*_3,*S*_ ∩ *X*_4,*W*_ and *F*_*Y*_ = *Y*_2,*S*_ ∩ *Y*_3,*W*_ ∩ *Y*_4,*S*_. If we presume that for each epoch and each synapse population, synapses are assigned randomly to the strong or weak population, then the expected values of *sF*_*X*_ and *sF*_*Y*_ are respectively 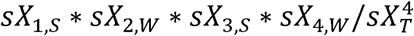 and 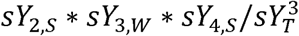. In Table 4, the expected and actual sizes of these fluctuating populations are respectively reported in columns E and A.

### Latent learning demonstration

To create a learning system that exhibits reward-dependent mapping of inputs to action choices, the baseline modeling approach is altered as follows. To simplify the development of input-choice mapping, we restrict the input set to the eight test stimuli, in input configuration 1, and during each stimulus period randomly select just one for presentation. To enable the network’s exhibition of choice behavior, we add to our system four “choice” neurons, such that the first of these activated subsequent to the presentation of a stimulus is regarded as the network’s choice. To enable the learning of specific input-choice preferences, we connect every output neuron to every choice neuron, augment the active synapses when the network’s choice is the rewarded choice, and decrement the active synapses otherwise. The weights of the output-choice synapses are bounded by *WC*_*min*_ and *WC*_*max*_. To hasten learning, we induce the network to make choices (and therefore receive environmental feedback) by randomly selecting (for each stimulus period) a choice neuron for stimulation with a bias current. These various model elements are captured Eq. 9 and Eq. 10, which are the respective update rules for choice neuron voltages and output-choice synapses. In these equations, *k, dwc*, 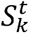, and *R*^*t*^ respectively correspond to the index of the choice neuron, a scaling factor of synaptic change magnitude, the existence of a choice neuron bias current, and the presence of a reward signal. Note that during simulation segments corresponding to latent learning, all 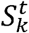 and R^t^ are fixed at 0, while for periods of active learning, 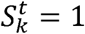 for only one value of *k* at any particular time *t*, and *R*^*t*^ = 1 (0 otherwise) if and only if the input and choice correspond to one of the simulation’s rewarded input-output pairs.

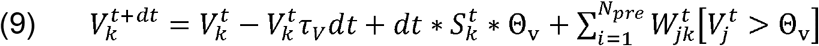

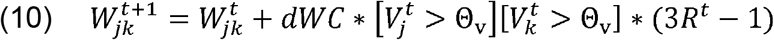

### Materials & statistical analyses

All simulations and data analyses are performed in Matlab (Natick, MA). All p-values above. 001 are reported exactly. For all statistical tests performed, a p-value below. 05 was considered significant. We use t-tests (matched pair, when applicable) to analyze distributions for which Lilliefor’s test does not suggest non-normality (i.e. for which p>.05); otherwise, we use rank sum tests. The n values reported for rank sum tests correspond to the size of each population considered in the comparison; unless otherwise noted, both populations are of the same size.

### Parameter values

Parameter values used in the models described above are provided in the table below. The objective that guided our choices was that we wanted our model to exhibit certain learning and connectivity development outcomes. Our chosen parameter values, and small variations thereon, generally achieve these objectives, but larger changes to individual parameters result in qualitatively different outcomes; for example, a shortened (lengthened) candidate synapse lifetime *µ* results in a very sparse (dense) arbor of stabilized synapses. Changes to multiple parameters in tandem, for their part, can sometimes result in strikingly large or small changes to system behavior. We saw (Figure 1E), for example, that very little changes when our generally-used values for *µ* and Θ_h_ (threshold for reinforcement-dependent stabilization) are both halved, because in this case two adversarial processes (candidate collapse vs. stabilization) are affected equally. These notions of process interaction and qualitative behavior preservation are what motivate our rare, but occasionally made, manual parameter value adjustments between experiments. For example, two of our modifications to the baseline model, introducing PING and simplifying the input module, both reduce the net input to neurons in the output module; a concurrent reduction to *r*_*o*_ (output neuron activity set point) helps preserve the manifestations of Hebbian competition (e.g. anterograde interference).

Table of simulation parameters

| Parameter | Description | Value |
| --- | --- | --- |
| $dt$ | Size of time discretization relative to a stimulus period | .01 |
| $S$ | Magnitude of bar stimulus | 270 |
| $\tau_v$ | Time constant, voltage decay | .5 |
| $\Theta_v$ | Threshold, voltage at firing (resting is 0) | 50 |
| $r_o$ | Set point, output neuron activity level | 20 (8, 2) |
| $\tau_F$ | Time constant, input multiplier change | .01 |
| $\tau_r$ | Time constant, output neuron activity level decay | .06 |
| $q$ | Firing time separation ceiling for Hebbian reinforcement | .1 |
| $L$ | Magnitude of activity-dependent synaptic weight decrement | .2 (.16) |
| $G_r$ | Strength of LASG | 4 (6, 0, 8) |
| $W_{min} (WC_{min})$ | Minimum synaptic weight, input-output (output-choice) synapses | 2 (1) |
| $W_{max} (WC_{max})$ | Maximum synaptic weight, input-output (output-choice) synapses | 10 (5) |
| $\Theta_g$ | Threshold, candidate integrator at generation | 8 (1.4) |
| $\mu^I$ | Initial value, collapse countdown timer | 40 |
| $\Theta_h$ | Threshold, total Hebbian reinforcement at stabilization | 24 (19.2) |
| $W_{OP}$ | Output-to-inhibitory synaptic weight (PING experiments) | 4 |
| $P(active)$ | Input neuron activation probability (linearized model) | .07 |
| $dWC$ | Scaling factor, reward-modulated synaptic adjustments (latent learning demonstration) | .2 |

Parameters are listed in the order introduced in this section. In the “Value” column, the first value provided is the value most commonly used. For $r_o$ , $L$ , $G_r$ , and $\Theta_h$ , the first (or only) value within parentheses is used during experiments with PING. The second parenthetical value for $r_o$ is used in experiments with the linearized model. The second and third parenthetical values for $G_r$ are respectively used during no-LASG experiments and the latent learning demonstration. The parenthetical value for $\Theta_g$ is used in no-LASG, high-candidate experiments.

